# Bilateral Intracranial Beta Activity during Forced and Spontaneous Movements in a Hemi-PD Rat Model

**DOI:** 10.1101/2020.08.16.252957

**Authors:** Soheil Mottaghi, Sandra Kohl, Dirk Biemann, Samuel Liebana, Ruth Montano, Oliver Buchholz, Mareike Wilson, Carolin Klaus, Michelle Uchenik, Christian Münkel, Robert Schmidt, Ulrich G. Hofmann

## Abstract

Cortico-basal ganglia beta oscillations (13-30Hz) are assumed to be involved in motor impairments in Parkinson’s Disease (PD), especially in bradykinesia and rigidity. Various studies have utilized the unilateral 6-OHDA rat PD model to further investigate PD and test novel treatments. However, a detailed behavioral and electrophysiological characterization of the model, including analyses of popular PD treatments such as DBS, has not been documented in the literature. We hence challenged the 6-OHDA rat PD model with a series of experiments (i.e. cylinder test, open field test and rotarod test) aimed at assessing the motor impairments, analyzing the effects of Deep Brain Stimulation (DBS), and identifying under which conditions excessive beta oscillations occur. We found that hemi-PD rats presented an impaired performance in all experiments compared to the sham group, and DBS could improve their overall performance. Across all the experiments and behaviors, the power in the high beta band was observed to be an important biomarker for PD as it showed differences between healthy and lesioned hemispheres and between PD and sham rats. This all shows that the 6-OHDA PD model accurately represents many of the motor and electrophysiological symptoms of PD and makes it a useful tool for the pre-clinical testing of new treatments and further investigations into this disease.

## I. Introduction

Parkinson’s disease (PD) is a neurodegenerative disorder which affects an estimated 10 million patients worldwide (Dorsey et al. 2018). The disease is famously characterized by both motor and nonmotor symptoms, including decreased and inhibited movements, resting tremor, rigidity, sleep issues, cognitive dysfunction, and depression (DeMaagd & Philip, 2015). It has been shown that neural loss in the nigral dopaminergic inputs to the striatum is one of the main causes of the condition. This leads to a major alteration in the neural activities of the cortico-basal ganglia loop (CBGL), which adversely affects the ability to make voluntary movements (Brown et al., 2001; Kühn et al., 2005; Levy et al., 2002). In the presence of normal dopaminergic drive, the activity of CBGL neurons is largely desynchronized. However, upon the loss of dopaminergic neurons, in idiopathic PD and experimental models of the disease, neurons of the subthalamic nucleus (STN), (internal and external) globus pallidus (GP) and substantia nigra pars reticulata (SNr), lose their independence and show increases in burst firing and synchronization of activity (Filion and Tremblay, 1991; Nini et al., 1995).

The synchronized neural oscillations in the aforementioned areas mostly have frequencies within the beta band (13–30 Hz), a frequency band which has been shown to correlate with bradykinesia and rigidity in PD patients (Kuhn et al., 2008). This causes the spectral power of the beta band to be higher than that of other frequency bands (e.g. alpha, gamma, etc.) in PD patients. The remaining spectral power of the synchronized oscillations is typically in the theta range (6-12Hz) and appears to be phase-related to the symptom of resting tremor. To confirm the causal relation between motor impairments and the excessive synchronization of beta band basal ganglia neural activity, Eusebio et al. (2008) investigated the effect of direct low frequency STN-DBS and reassuringly showed that it leads to a worsening of bradykinesia symptoms in human PD patients.

It hence comes as no surprise that research to-date suggests that common treatments for bradykinesia, such as levodopa medication or high-frequency deep brain stimulation (DBS), work by suppressing the synchronized beta band oscillations in the cortico-thalamo-basal ganglia circuit (Delaville, McCoy, Gerber, Cruz, & Walters, 2015; Dorval, Kuncel, Birdno, Turner, & Grill, 2010; Kühn et al., 2005; Levy et al., 2002). This suppression is predominantly observed in the STN (Gatev, Darbin, & Wichmann, 2006; Plenz & Kital, 1999; Stein & Bar-Gad, 2013) and the motor cortex (Yamawaki, Stanford, Hall, & Woodhall, 2008) of PD patients. It is, however, important to note that this suppression of neural activity in the cortico-thalamo-basal ganglia circuit only leads to motor improvements in patients who already present deteriorated capabilities, and will instead impair the performance of PD patients whose motor capabilities are within normal limits (Chen et al., 2006).

Adaptive deep brain stimulation (aDBS) is emerging as a promising enhanced treatment for PD which overcomes several limitations of conventional DBS (Little et al., 2020). Since physiological biomarkers derived from CBGL beta band oscillations are currently the front-runners as control signals for aDBS (Castaño-Candamil, Mottaghi, Coenen, Hofmann, & Tangermann, 2017; Little et al., 2013, 2014), the development of future control algorithms for this treatment (Hoang, Cassar, Grill, & Turner, 2017; Neumann et al., 2019) will surely benefit from a framework for the systematic testing of these biomarkers using accessible and established animal models of PD.

In this study we present an investigation of the suitability of the unilateral 6-hydroxydopamine (6-OHDA) hemi-PD rat model (Ungerstedt, 1968) as a framework for PD pre-clinical research. The unilateral 6-OHDA rat model simulates PD by unilaterally injecting the highly specific neurotoxin 6-hydroxydopamine (6-OHDA) into either the medial forebrain bundle (MFB) or the substantia nigra, causing substantial ipsilateral dopamine loss (Henderson et al., 2003; Ungerstedt, 1968). This creates what is known as a hemi-parkinsonian (hemi-PD) rat, where one hemisphere of the rat brain is significantly damaged, and the contralateral side to the lesion serves as an in-animal control allowing for behavioral comparisons within the rat (Lundblad et al., 2005).

Our work aims to characterize the motor impairments displayed by the unilateral 6-OHDA model, as well as the conditions under which an excess in the spectral power of the beta frequency band is exhibited in the neural oscillations of the primary motor cortex (M1) and the subthalamic nucleus (STN) of animals lesioned according to this model. Since the hemi-PD model only simulates the dopamine-loss aspect of PD, we thought that investigating behavior-related changes in beta power could provide interesting insights. To achieve this, a series of tests assessing the different movement capabilities of the animals were performed, including the cylinder, open field and rotarod tests. In addition to the characterization of the model’s effect on animal motor capabilities, the effect of standard DBS on the improvement of motor impairments and the normalization of beta oscillations was investigated through the above tests as well. Finally, all throughout our study we assessed the suitability of beta power in the STN as a biomarker to control stimulation in aDBS.

In a similar study of the 6-OHDA model, Degos et al. reassuringly observed PD’s characteristic excessive synchronization of beta band oscillations in the motor cortex of PD rats, however, the bradykinetic/akinetic symptoms of their model appeared before this synchronization was detected (Degos, Deniau, Chavez, & Maurice, 2009). Additionally, their study observed that this enhanced beta power is only significant while the animal is awake and not during periods of sleep. The recent work published by Swan et al. on the causal relationship between STN beta band oscillations and bradykinesia/akinesia symptoms in hemi-PD rats has further shown that applying STN-DBS at beta band frequencies does not have a significant impact on the motor performance of hemi-PD rats, leading the authors to claim that beta oscillations in the STN are an insufficient biomarker for motor symptoms in this model (Swan, Schulte, Brocker, & Grill, 2019).

We expand on both these studies by not only confirming some of their findings, but also providing an in-depth description of the motor impairments that the 6-OHDA model causes in hemi-PD rats and how these compare to the symptoms in human PD. We also evaluate the degree to which standard STN-DBS ameliorates the lesioned animal’s motor impairments, something which has not been done in as much detail as in our study before. Throughout all the experiments we continuously record electrophysiological signals from the STN and M1 in order to characterize the changes in the neural activity brought about both by the model and by the several activities that the animals perform, having as one of our main aims that of assessing the suitability of STN beta power as a biomarker for aDBS - something which has been contested in the recent literature as mentioned above (Swan, Schulte, Brocker, & Grill, 2019).

The main findings of our experiments were that the 6-OHDA PD model accurately simulates the motor and balance impairments of PD in both free and constrained motion and that DBS is capable of restoring the capabilities of the animals back to healthy levels. We also discovered that excessive beta power is prominent in PD rats in all experimental settings i.e. in free and forced motion at all speeds – except when the animal was static. An analysis of the correlation between speed and spectral power further revealed significant correlations only in forced movement and was positive in the beta band and negative in the theta band. We also noticed that excessive beta power was more prominent in M1 than in the STN, especially during movement initiation.

We hope that through this work, the better understanding of the 6-OHDA PD rat model that we provide will enable the model to be used more frequently as a framework for PD pre-clinical research. The possibility of using an animal model to accurately investigate the efficacy of new and alternative treatments for PD will surely accelerate and facilitate current research, ultimately leading to the improved quality of life of many PD patients.

## II. Materials and Methods

All animal procedures were conducted in conformity with relevant institutional rules in compliance with the guidelines of the German Council on Animal Protection. Protocols were approved by the Animal Care Committee of the University of Freiburg under the supervision of the Regierungspräsidium Freiburg (approval G15/031) in accordance with the guidelines of the European Union Directive 2010/63/UE.

### A. Surgical Procedure

Two groups of adult female Sprague-Dawley rats (290-310g), consisting of 13 hemi-parkinsonian (PD-group) and 12 non-lesioned rats (sham-group), were used for this study. All animals were acquired from Charles River Laboratories (Germany) and were housed under temperature-controlled conditions in a 12-h light-dark cycle, with access to water and food *ad libitum*. Rats were allowed to acclimate to these conditions for a minimum of two weeks prior to any experimental procedures in order to reduce unnecessary stress.

Prior to surgery, all rats underwent at least 7 days of handling to familiarize them with the experimenter. During each surgery, the rats were anesthetized with oxygen (0.15 l/min) and isoflurane (Abbvie, USA). Anesthesia was induced with 4% isoflurane and gradually lowered to 1.5% after placing the animal into the stereotaxic frame (David Kopf, USA). Breathing, reflexes and depth of anesthesia were monitored throughout the duration of the surgery.

All PD-group animals underwent two consecutive stereotaxic surgeries, with a two-week recovery time between each surgery. In the first, the 6-OHDA was injected, and in the second, custom-made microelectrodes for stimulation and recording were implanted. During each surgery, holes were drilled at the injection or implantation site, and the dura was resected using a fine needle. The injection canula or electrode was subsequently lowered manually at a rate of approximately 200µm/second. Four minutes were allowed to pass after each injection, with the needle still inserted, to allow the 6-OHDA to adequately diffuse into the target. After electrode implantation, the skull aperture around the implanted electrode was filled with bone wax. Once in place, electrodes were fixed to a nearby stainless-steel screw anchor (0-80 x 1/8; Plastics One) using a 2-compound dental cement (Palapress; Heraeus Holding GmbH; Germany).

Sham animals were only subjected to the electrode implantation surgery, which followed a similar procedure to that of the PD group (see Electrode Implantation for more details).

#### i. 6-OHDA Lesion and Apomorphine Test

Animals assigned to the PD group were unilaterally lesioned with a 6-OHDA solution (3.6mg 6-OHDA, 20mg ascorbic acid, 10ml 0.9% NaCl) injected into the right medial forebrain bundle. Table 1.1 contains the lesion coordinates, 6-OHDA volume and the injection rate. The Apomorphine rotation test was performed after a recovery period of 2 weeks to test dopamine depletion intensity. Animals were deemed sufficiently lesioned if at least 3 contralateral (to the lesion) rotations on average were observed per minute for 30 minutes after the subcutaneous injection of a 0.1ml/100 gr. Apomorphine solution (1mg Apomorphine, 2mg ascorbic acid acquired from Sigma-Aldrich Chemie GmbH, Germany and 20ml NaCl).

**Table 1.1.** Lesion coordinates in MFB, quantity of 6-OHDA per coordinate and injection rate.

|  | AP (mm) | ML (mm) | DV (mm) | Quantity | Injec. Rate |
| --- | --- | --- | --- | --- | --- |
| MFB | -4.4 | -1.2 | -7.8 | 2.5 μl | 1 μl/min |
| | -4.0 | -0.8 | -7.2 | 3.0 μl | 1 μl/min |

#### ii. Electrode Implantation

All rats received a bilateral implantation of Platinum-Iridium (70% Pt, 30% Ir) bipolar recording electrodes with 10µm and 75µm tip diameter and separation respectively (Science Products GmbH, Germany) into the primary motor cortex (M1) and STN, to achieve a total of eight recording sites for each rat brain. The coordinates of the implants are available in Table 2. Additionally, rats in the PD group received custom-made bipolar stimulation electrodes, made of intertwined 50µm Platinum-Iridium (70% Pt, 30% Ir) microwires (Science Products GmbH, Germany). The stimulation electrodes were implanted into the STN ipsilateral to the lesion and adjacent to the recording electrodes.

**Table 2.** Electrode implantation coordinates in M1 and STN

|  | AP (mm) | ML (mm) | DV (mm) |
| --- | --- | --- | --- |
| M1 | +2.5 | ±3.0 | -1.6 |
| STN | -3.8 | ± 2.4 | 8.0 |

### B. Animal Experiments

In all experiments, video recordings of animal behavior and electrophysiological recordings of brain signals were collected. Electrophysiological data in the form of local field potentials (LFP) and spiking data (SPK) was recorded from the bilaterally-implanted recording electrodes in the STN and M1. Data was collected using a 32-channel wireless head stage (Multi-channel System GmbH, Germany) and an AlphaLab SnR system (AlphaOmega, Israel). Prior to all behavioral tests, an adequate stimulation strength for each animal of the PD group was individually determined by titrating the current amplitude such that explorative behavior was observed, but stimulation-related side effects were not. In this way, we compensated for possible variations in electrode placement and shape. Stimulation was applied at a frequency of 130Hz using a biphasic rectangular pulse and a pulse width of 65µs in each of the following behavioral tests.

#### i. Cylinder Test

The cylinder test measures spontaneous forelimb use, body movement and exploratory activities (Cenci & Lundblad, 2005). Animals were not introduced to the cylinder prior to the experiment in order to test their motivation and capability to explore a novel environment.

Each rat was placed in a transparent cylinder (acrylic glass, 19 cm diam., 35 cm ht.) for a total of two minutes (see Figure 1.A-B). A Logitech C920 HD video camera (Logitech, Switzerland) was used to record the animals from a ventral viewpoint (30fps). Rats belonging to the PD group (n=13) were split into two groups, those receiving DBS during the experiment (PD-DBS ON, n=7) and those which did not (PD-DBS OFF, n=6). Sham rats (n=12) never received DBS. The experimental schedule is shown in Figure 1.C.

**Figure 1.**
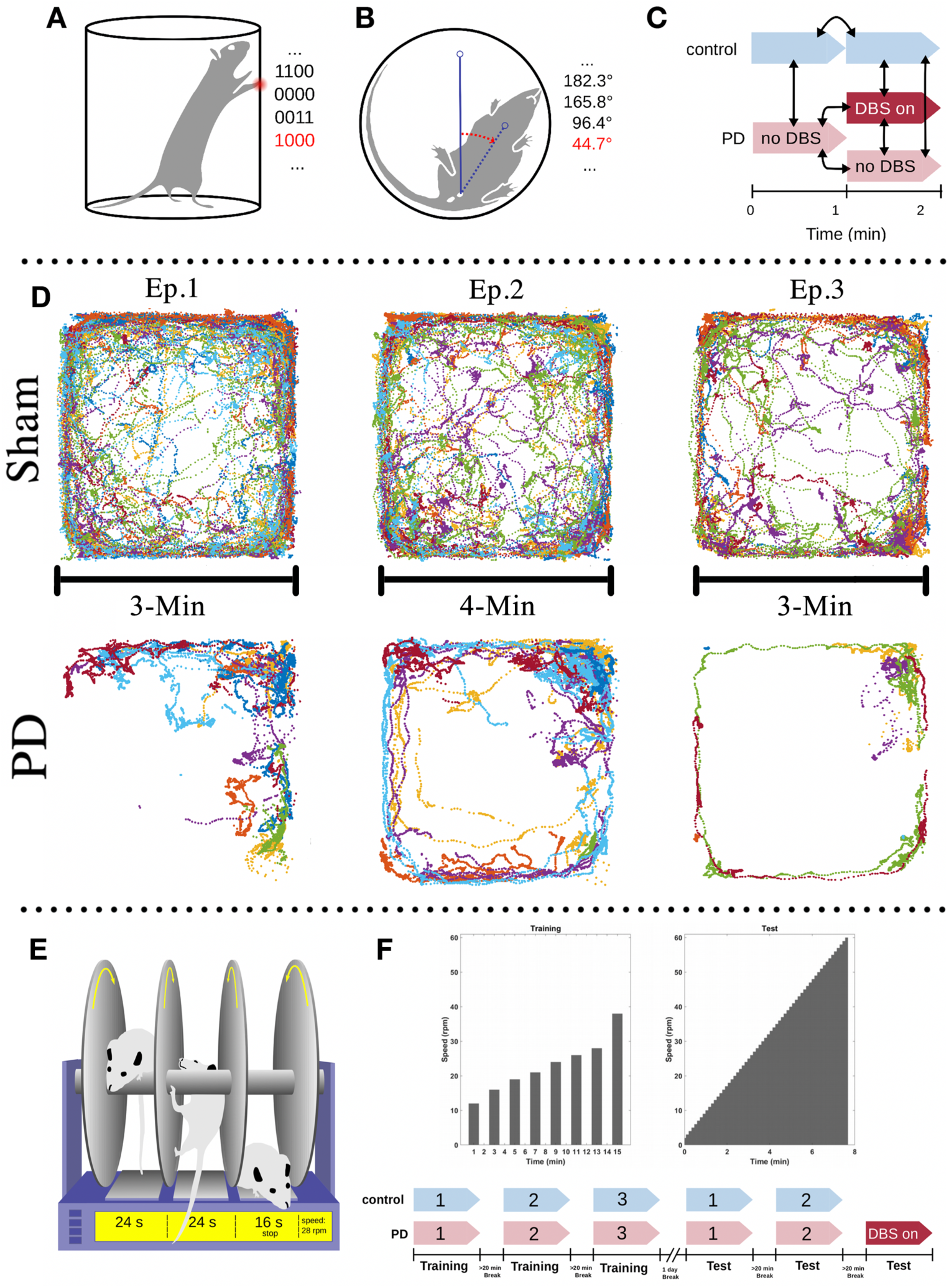
Three experimental paradigms were investigated in this study to challenge the animals’ motor capabilities and find the conditions under which excessive beta synchronization is observed. A-B: The position of the forepaws and the body angle of the animals were measured (e.g. 1100 = both front paws on wall, 0000 = rearing with no support, 0011 = both front paws on floor, 1000 = left paw on wall and right paw not in contact). C: The animals were monitored for two minutes. A subgroup of PD animals received DBS treatment at the onset of the second minute, while the rest of the PD group remained untreated. D: Open field area, a 74cm x 74cm square box, with the exploratory activities of PD and Sham animals during 10 minutes superimposed. Note how sham rats explore most of the area available to them (with a tendency to avoid the center which is a natural reaction for rats), whereas PD rats remain on the corners and explore much less than sham rats. Notice also the significant difference in the trajectories between stimulated PD rats (episode 2) and non-stimulated PD rats (episode 1 and 3). This shows that DBS considerably increases the rats’ motivation to explore and corrects major motor impairments. E-F: All the animals were trained on the rotarod one day prior to the test day. On the test day, both the sham and PD group performed two sessions of testing without DBS. A subgroup of the PD rats (n=7) performed a third round while receiving DBS treatment. The other subgroup performed the third round without DBS (n=6).

The observed behavior of all rats was classified into three distinct patterns: rearing, stepping and inactive behavior. During rearing, rats stand on their hind limbs while their forelimbs touch the cylinder wall to maintain balance (Figure 1.A). Stepping was defined as episodes where the orientation of the animal’s body shifted >45 degrees while its forepaws alternated between touching the ground and being held in the air (Figure 1.B). Any behavior that could not be defined as either rearing or stepping was classified as inactive.

To quantitatively assess the amount of time spent in each of the three behavioral categories, the distinct positions of the rat’s forelimbs were identified and converted into a four-digit binary code, updated every 0.25s. The first two digits represent the left and right forelimb respectively and indicate whether the paws touch the cylinder wall (1) or not (0). Similarly, the last two digits show whether the forelimb paws touch the ground (e.g. 1100 = both front paws on wall, 0011 = both front paws on floor, 0000 = rearing with no support, 0110 = right paw on wall and left paw on floor). The body rotation was also measured (in θ/s) using Tracker video analysis software (©2016, Douglas Brown, https://physlets.org/tracker) combined with a virtual protractor overlay on the video, with the protractor vertex centered on the center of the urinary meatus of the rat. The endpoint of one protractor leg was aligned with the upper chest of the animal while the remaining leg served as a reference for rotation. After collection, the binary forelimb states and rotation measurements were subsequently analyzed in MATLAB (MathWorks, USA).

#### ii. Open Field Test

To measure the animal’s locomotor performance and explorative behavior in a novel environment, an open field test (Seibenhener et al., 2015) was performed. The rats were placed in a featureless square chamber (74 × 74 × 30 cm) and left free to explore for 10 minutes while their location was tracked using BioObserve Viewer II software (BioObserve GmbH, Germany, 25fps). The experimental layout, together with the superimposed trajectories of the tested animals, are depicted in Figure 1.D. The experiment was divided into three episodes. During the first 3 minutes (episode 1), no rat (n=25) received DBS stimulation. Next, PD-group animals were divided into two subgroups; one received DBS for 4 minutes (PD-DBS ON, n=7) while the other did not (PD-DBS OFF, n=6). In the last episode, DBS was turned off for all the PD rats (n=13) and stayed off for the remaining 3 minutes. Sham rats (n=12) were left free to explore with no DBS for the entirety of the experiment. The environment was cleaned after each trial to prevent any lingering olfactory signals from interfering with behavior. As in the cylinder test, electrophysiological and video tracking data was collected and subsequently analyzed offline.

#### iii. Rotarod

The rotarod test, first described by Dunham & Miya (1957), is widely used to assess the effect of brain injuries or experimental drugs on motor function in rodents (Cartmell et al., 1991; Bohlen et al., 2009). No experienced observer or nominal scoring procedure is required, as the test yields a discretely measurable variable (time or speed) which can be used to quantify motor behavior in an objective manner. To perform this test, the rat is placed on a rotating rod that mimics a treadmill. The rod is suspended high enough over the ground so that the rat is naturally motivated to avoid falling, but low enough to avoid any injuries should a fall occur. The rotational speed of the rod is then gradually increased until, when the rat eventually loses its grip or balance, it falls onto a switch plate beneath the rod. Both the speed of the rod and the time that the animals remained on the rotarod are measured and recorded (Figure 1.E).

The experimental schedule spans over two days: on the first day, all rats (n=25) were trained on a rotarod (Rat Rota-Rod 47700; UGO Basile S.R.L. Gemonio, Italy) in three sessions with 20 minutes between each session. The sessions consisted of 8 trials with different velocity settings of the rod (12, 16, 19, 21, 24, 26, 28 and 38 rpm) and a maximal duration of 60s. Each trial was followed by a resting period of one minute (Figure 1.F). On the second day, the rotarod was set to accelerate from 2 to 60 rpm with 1 rpm steps over the course of 7 min. 52 sec. All rats performed the test in two sessions with a break of at least 20 minutes in between the sessions. The PD rats were then split into two groups where PD-DBS ON rats (n=7) performed an additional third session while receiving DBS and PD-DBS OFF (n=6) did the same but without DBS.

### C. Electrophysiological Analysis

All of our data analysis was performed in MATLAB 2017a (Mathworks, USA) and Python (Python Software Foundation, CWI). LFP signals were resampled at 1.1 kHz. Power spectral density (PSD) was calculated using Thomson’s multitaper PSD estimate (5 Slepian tapers). In this paper, the right hemisphere of PD-group animals corresponds to the lesioned side, while the left hemisphere is the intact (unlesioned) side. Sham rats only had recording electrodes and hence both of their hemispheres were intact.

The Euclidean Distances (ED) between the averaged PSD’s of different brain regions were used to quantitatively compare electrophysiological behavior across the brain. In our experiments these were used to quantify the difference in beta band power (13-30Hz) between sham rats and PD rats as follows:

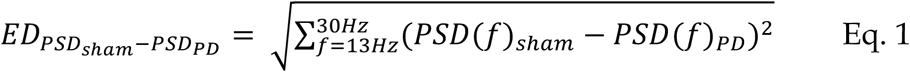

The behavioral results and normalized band powers were statistically analyzed using a nonparametric Wilcoxon rank-sum test, unless mentioned otherwise. Kendall’s rank correlation coefficient was used to assess the linear correlation between band power and speed.

### D. Euthanasia & Histology

After having finalized all experimental testing, chronically implanted rats were euthanized with an overdose of isoflurane and were perfused transcardially with a 4% formaldehyde solution (PFA in phosphate buffer). Their brains were removed, post-fixed in PFA for 7 days and stored in 30% sucrose, after which they were cut into coronal sections (40 µm) along the probe’s implantation trajectory with a cryostat (CryoStar NX70, ThermoFisher scientific, USA). Sections were collected on glass slides and stored at 4°C until further processing. Tyrosine Hydroxylase TH staining was then used as a marker to identify the presence (or lack of) dopaminergic neurons and thus assess the success of the lesion. The sections were also used to verify the correct positioning of the recording and stimulating electrodes.

## Results

Through our results, we have provided an in-depth characterization of the motor impairments caused by the unilateral 6-OHDA lesion of the hemi-PD rat model and have investigated the conditions under which excessive beta power can be observed in the STN and M1 of animals lesioned according to this model. The behavioral analysis for each experiment had as its aim that of comparing the impairments of the PD group with the motor abilities of the sham group. The electrophysiological analysis was then carried out according to the results of the behavioral assessments to investigate the differences in neural activity arising from the different behaviors. To this end, the recorded LFP signal (0.3-300Hz) was filtered into θ (6-12Hz), low (13-21Hz) and high (21-30Hz) β bands (see Figure 2).

**Figure 2.**
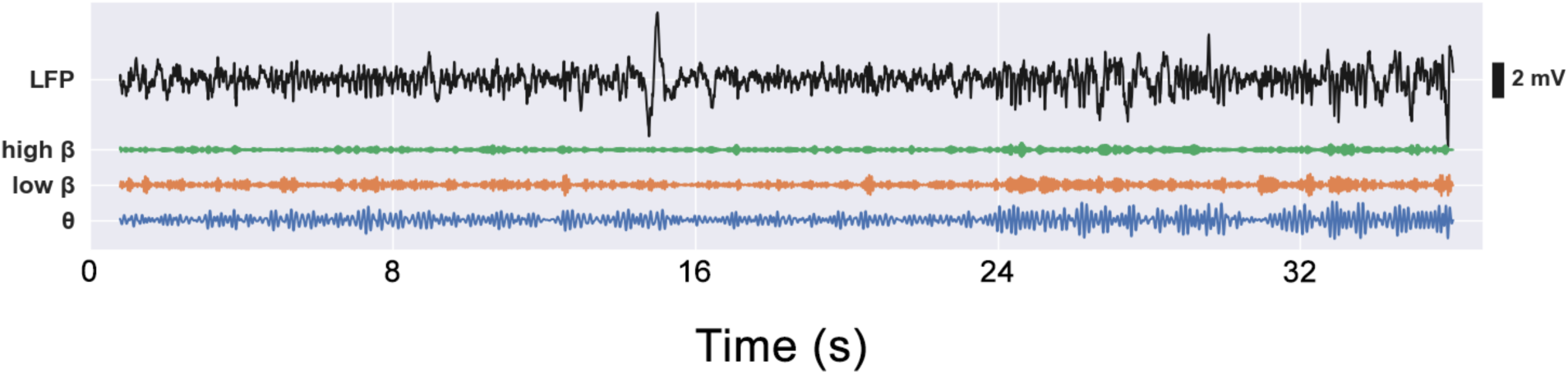
Electrophysiological LFP signals (0.3-300Hz), as well as video footage, were captured during each experiment simultaneously and were analyzed offline. LFP signals were filtered into θ (6-12 Hz), low β (13-21Hz) and high β (21-30Hz) frequency bands.

### A. DBS Restores Exploratory Movement Patterns in the 6-OHDA Hemi-PD Rat

The cylinder and open field (OF) tests investigated exploratory movement patterns in both the PD and sham animal groups, since the animals were free to move and explore in both environments. The cylinder environment favored rearing and other stationary activities while the OF challenged their locomotion capabilities.

In the cylinder test, the time that every animal spent in each of the 3 behavioral categories (i.e. rearing, stepping and inactive) was measured and analyzed individually for each category. To establish a baseline behavioral pattern, the performance of the sham rats during the first minute of the experiment was compared to their performance during the second minute; thus focusing on any behavioral changes resulting uniquely from the time spent in the cylinder. A significant decrease in the sham rats’ rearing time and a significant increase in the duration and frequency of their inactive episodes was observed from this comparison (p_rear_<0.05, p_inactive_<0.01). However, the difference in the time spent stepping was insignificant (see Figure 3.A). This shows that there is a natural tendency for the rats to explore less the longer they spend in the cylinder environment, as expected. These results are useful to gauge the scale of the decrease in exploratory behavior arising from a natural ‘boredom’ so that we can then compare it to the decrease in exploration caused by the 6-OHDA lesion.

**Figure 3.**
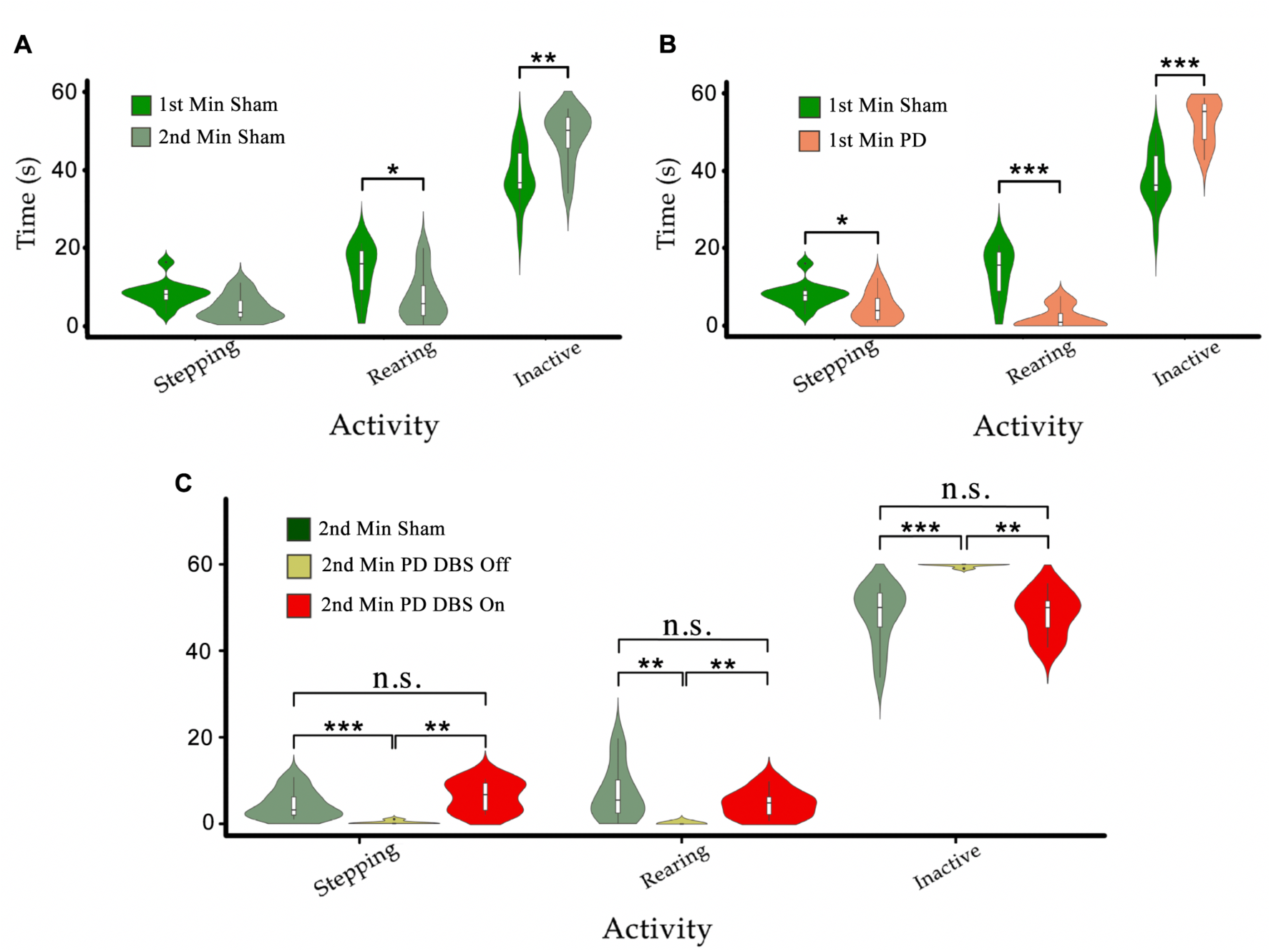
(A) The time sham animals spend in stepping, rearing and inactive conditions during the first minute of the tests was compared to the second minute. We observe decreases in stepping and rearing behavior and an increase in the time spent in the inactive state. (B) The comparison between the behavior of PD and sham rats during the first minute. We see a similar, yet more pronounced difference than in (A). (C) The behavioral patterns of PD-DBS ON, PD-DBS OFF and sham animals during the second minute of the cylinder test. We see a similar pattern for PD-DBS ON and sham rats and a drastically impaired pattern in PD-DBS OFF rats, which spend nearly all of their time in the inactive state. Level of significance: (* p<0.05,** p<0.01,*** p<0.001).

A similar comparison between the PD and sham groups over the first minute shows that PD rats spent significantly less time stepping and rearing, and instead spent more time in the inactive state (p_stepping_<0.05, p_rear_<0.001, p_inactive_<0.001) (Figure 3.B). It can thus be seen that the difference in the time spent in stepping behavior between the PD and sham rats is only slightly more pronounced than in the above-described comparison between sham rats across time. The differences in the time spent rearing and in the inactive state are however much more pronounced between PD and sham rats than between sham rats at different times.

To evaluate the effect of DBS on the behavior of PD rats, the PD-group was divided into two subgroups: one PD subgroup (n=7, PD-DBS ON) was exposed to STN-DBS at the onset of the second minute of the experiment, while the other subgroup remained untreated (n=6, PD-DBS OFF). The results indicate that PD-DBS ON rats exhibited a similar behavioral pattern to the sham group during the second minute, whereas PD-DBS OFF rats exhibited significantly less stepping (p_Sham-PD-DBSOFF_<0.001 and p_PD-DBSON – PD-DBSOFF_<0.01, respectively) and rearing (p_Sham-PD-DBSOFF_<0.01 and p_PD-DBSON – PD-DBSOFF_<0.01, respectively), and instead spent significantly longer periods of time in the inactive state (p_Sham-PD-DBSOFF_ < 0.001 and p_PD-DBSON - PD-DBSOFF_ < 0.01, respectively) compared to the sham and PD-DBS ON groups (see Figure 3.C).

The data from the OF test shows similar results to that of the cylinder test. This data was analyzed for all groups by dividing the tracking data into three episodes: (1) the first 3 minutes of the experiment where no stimulation was applied; (2) minutes 4 to 8, where only the PD-DBS ON subgroup (n=7) received DBS, and the rest of the groups continued untreated; (3) the last 3 minutes, where once again no DBS was applied to any of the groups. Subsequently, we compared the behavioral patterns exhibited in each of the episodes across groups.

Figure 1.D illustrates overlaid tracking data for sham and PD group animals in each episode. Similar to in the cylinder test, it can be seen that the sham rats’ exploration declined as time progressed. Figure 1.D also shows that sham rats explore significantly more than PD-group rats, where PD animals tend to move less and like to stick to the borders of the arena. Although DBS increased the extent of exploratory behavior of PD rats in episode 2, when it was removed during episode 3, PD animals returned to their mostly inactive behavior.

In order to quantify and statistically compare these behavioral differences, four parameters were considered: average velocity, average time showing large movements (LM) (speed > 4cm/s, t > 2s), total distance travelled and average time spent immobile (speed < 0.5cm/s, t > 2s). Sham rats showed a decrease in average velocity, average time spent in LM and distance travelled if we compare the results from the 1^st^ and 3^rd^ episodes. On the other hand, the average time sham rats spent immobile increased throughout this timespan. This confirms the observations described earlier.

During the 1^st^ and 3^rd^ episodes (where DBS was not applied to any animal), significant differences in all four parameters were seen between the sham and PD groups (p_speed_, p_LM_, p_tr.dist._, p_immob._ <0.001). In the 2^nd^ episode, no significant difference was observed between any of the measured parameters for sham rats and the PD-DBS-ON subgroup, however the PD-DBS-OFF subgroup exhibited significantly less exploratory behavior in comparison. Figure 4.A-D depicts these behavioral results from the OF test.

**Figure 4.**
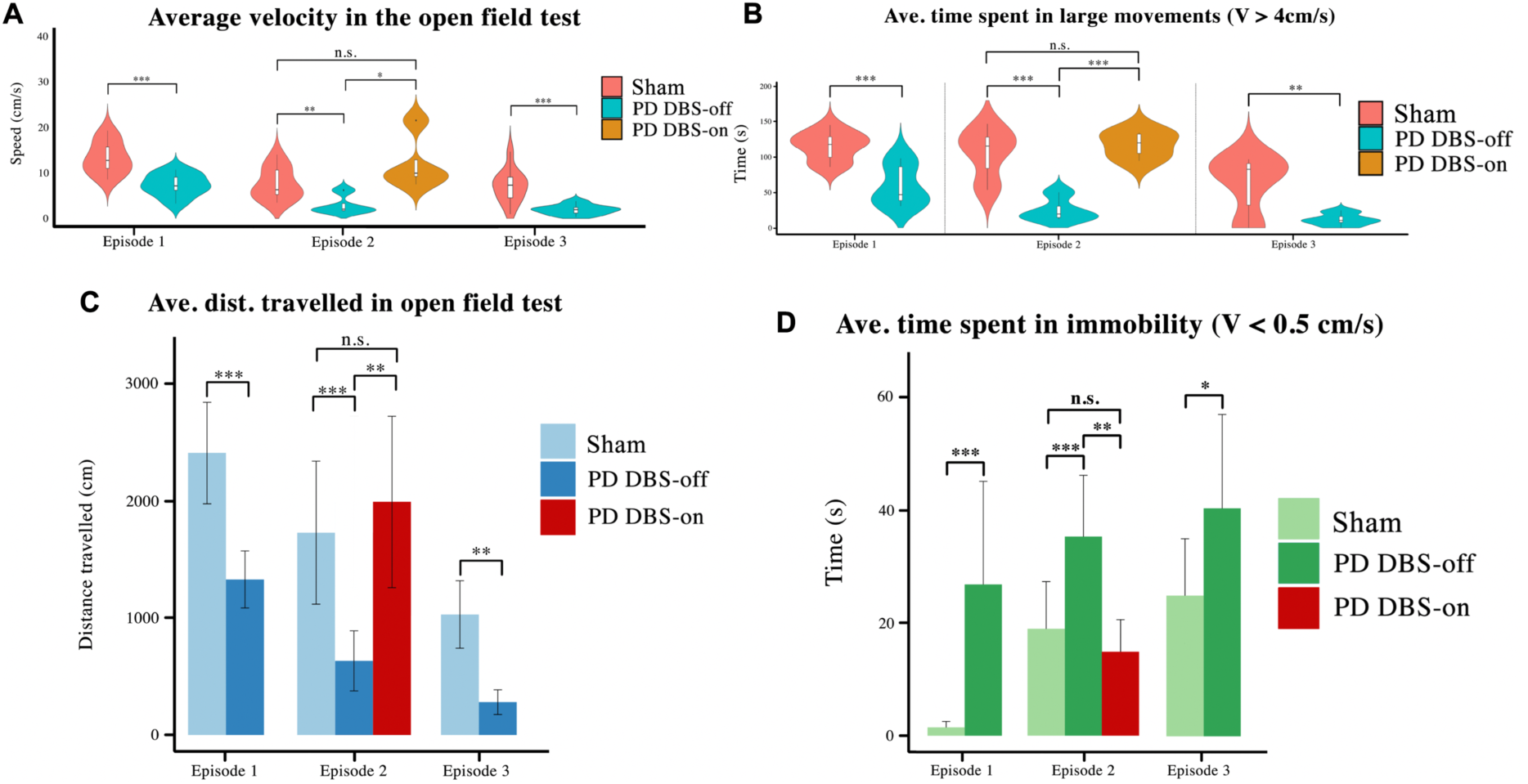
The behavioral features of the OF test were analyzed using the animals’ tracking data. (A-D) The average velocity, time spent in large movements, distance travelled and the time spent in immobility were selected as variables for comparison across groups. It is shown here that untreated PD animals exhibit lower velocity, distance travelled and time spent in large movements than treated and healthy animals, and instead spend more time in immobility. However, the PD subgroup that received DBS exhibits significant exploratory improvements and a significant reduction in their immobility spells. Level of significance: (* p<0.05, ** p<0.01, *** p<0.001).

The behavioral results from both the cylinder and OF tests hence show that we can reestablish severely impaired movement capabilities to close-to-healthy levels through the use of STN-DBS on subjects of the hemi-PD rat model.

### B. Distinct Movement-Dependent PSD Profiles in Hemi-PD Rats

The digitized behavioral patterns of the animals’ forelimbs in the cylinder test were synchronized with the time-frequency power analysis of the electrophysiological recordings in order to identify possible movement-dependent patterns. A digitized value > 4 (in its decimal representation) is considered a rearing position since it means that at least one of the animal’s forelimb paws is touching the wall.

Figure 5 shows spectrograms from an electrophysiological recording of both the unlesioned and lesioned M1 of one sample rat from the PD group, synchronized with a plot showing the rat’s progression through different forelimb positions. The lesioned hemisphere in the PD rat shows a strong increase in beta power (13-30Hz) compared to its unlesioned (intact) hemisphere in moments when the rat was about to enter the rearing position. This confirms the findings of (Degos, Deniau, Chavez, & Maurice, 2009) who also observed an excess in beta band power in PD rats of the unilateral 6-OHDA model. The correspondence of the peak in beta power with rearing episodes shows that the PD model is affecting neural activity during movement planning and initiation, which could explain the tendency of rats to remain in the inactive state as described earlier.

**Figure 5.**
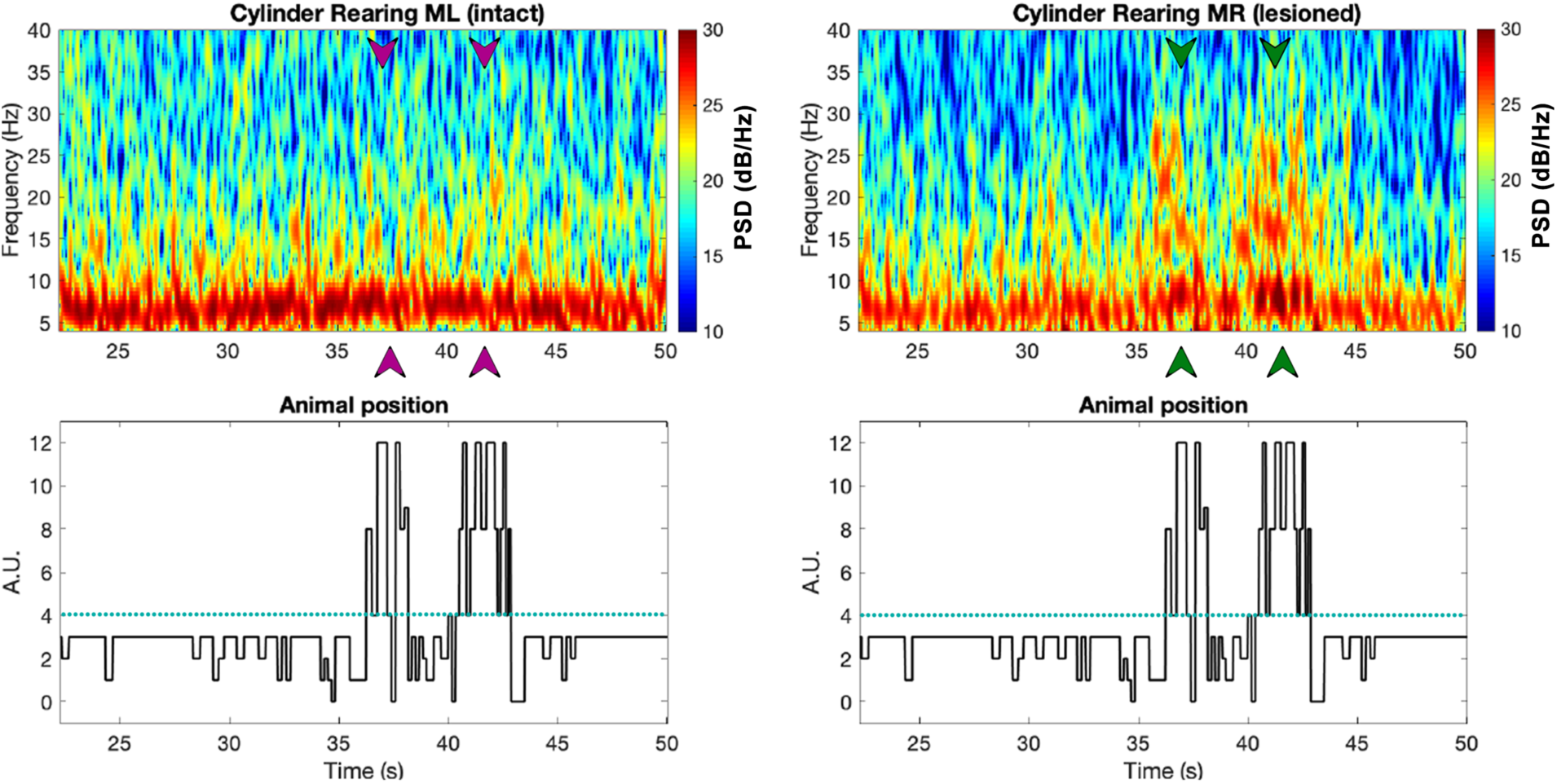
The top two plots show spectrogram examples from the intact (left) vs. lesioned (right) hemispheres captured from the M1 of a PD rat (R21), with the digitized behavioral pattern shown below each spectrogram. Note that the arrows in the spectrograms point at the times in which the animal was rearing, as can be seen from the forelimb position plot below (green and purple correspond to the lesioned and intact hemispheres respectively).

For both sham and PD rats, we also calculated the average band power within the frequency bands of interest (i.e. theta (6-12 Hz), low beta (13-21 Hz) and high beta (21-30 Hz)) for signals corresponding to the three behavioral patterns of the cylinder test (Figure 66). The results show distinct low and high beta band power differences between intact and lesioned hemispheres during rearing, significant differences only in low beta band power during stepping and no significant differences at any of the three frequency bands during inactive episodes. As a control, we also looked at the differences in the band powers between the two intact hemispheres of sham rats, finding no significant differences. This hence confirms that the differences in band powers for PD rats were due to the lesion.

These results show us that the differences in the spectra of electrophysiological recordings from the left and right M1 of PD and sham rats change depending on the behavior of the rats. It is clear from the above plots that in active episodes, the spectra of intact and lesioned hemispheres differ more than in no-movement episodes. This highlights the correlation of altered power spectra with motor symptoms in PD. Another important observation is that the differences in the spectra of lesioned and intact hemispheres are always concentrated around the beta band (both low and high), hinting at the beta band’s importance as a biomarker to indicate the presence of PD symptoms.

**Figure 6.**
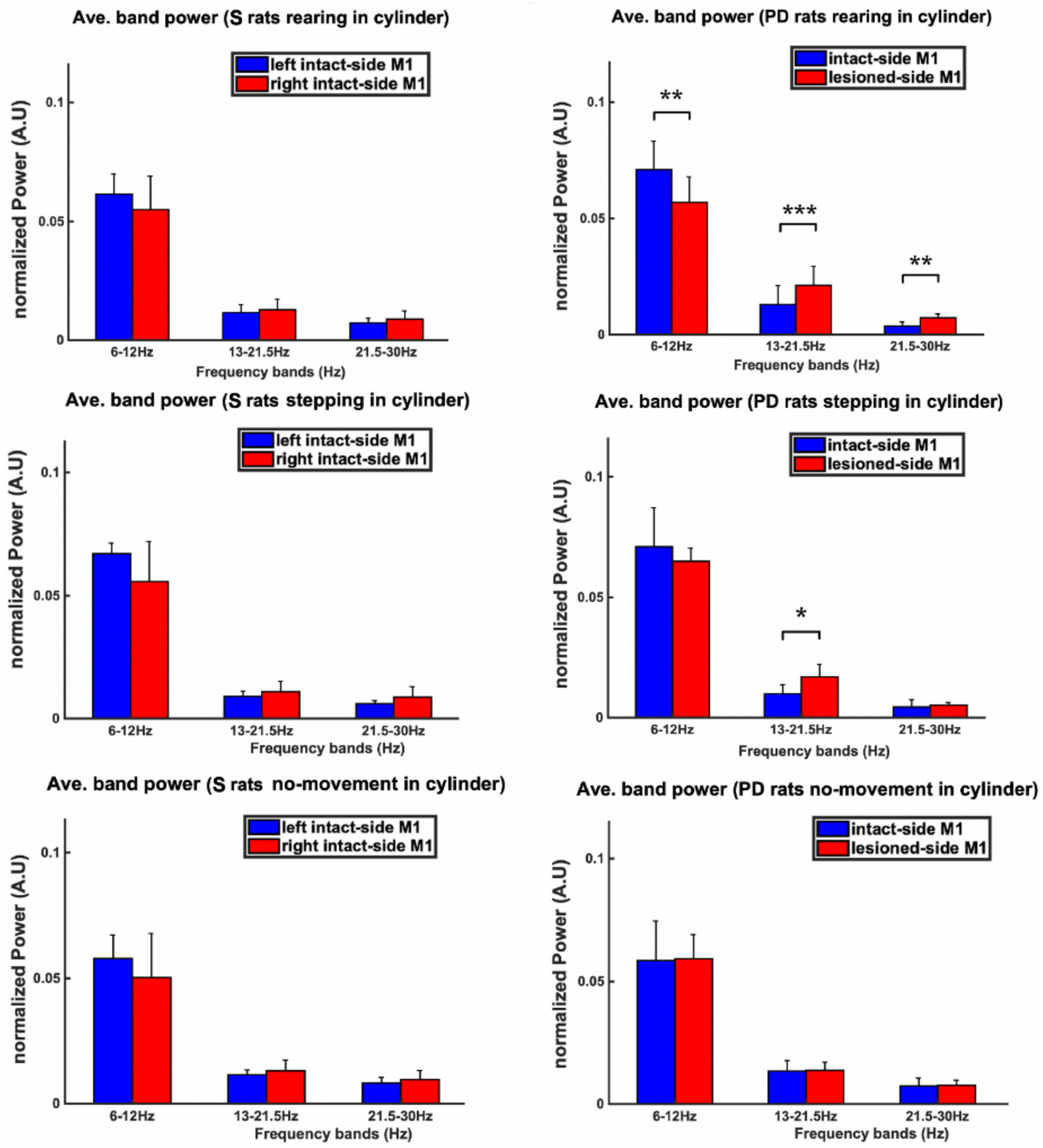
The power spectral density (PSD) during the three behavioral patterns of the cylinder test (i.e. rearing, stepping and no-movement) was analyzed. The PSD was averaged over theta, low beta and high beta bands measured from both hemispheres, resulting in three bins, one for each band. The averaged-PSD comparison of the right vs. left M1 of sham rats is shown in the left column, whilst the same comparison for PD rats is shown in the right column. It can be seen that during rearing in PD rats, both the low and high beta powers from the lesioned hemisphere (right hemisphere) are significantly higher than in the intact hemisphere. During stepping episodes, however, only the power in the low beta band is significantly higher in the lesioned than in the intact hemisphere. No significant differences between hemispheres can be observed for any band during inactive episodes in PD rats. Similarly, no significant differences are found by comparing the power in PD animals to that of sham group animals for inactive episodes. Level of significance: (* p<0.05, ** p<0.01, *** p<0.001).

### C. Excessive Beta Power Is More Prominent in M1 than in the STN

The prominence of excessive beta power in the lesioned hemisphere of PD rats was compared between recordings from M1 and the STN of PD and sham rats performing the cylinder test. To this end, we computed the PSD of M1 and STN recordings for all animals during rearing and stepping episodes - where the difference in beta power between intact and lesioned hemispheres had been found to be significant.

Figure 7 illustrates the PSD analysis of recordings from the STN and M1 of the lesioned hemisphere of PD rats during DBS-OFF and ON episodes, as well as from the corresponding intact regions of the sham group. In order to quantify the difference between the PSD’s from these regions, the Euclidean Distances (ED) between the averaged PSD’s were calculated as in Eq.1.

**Figure 7.**
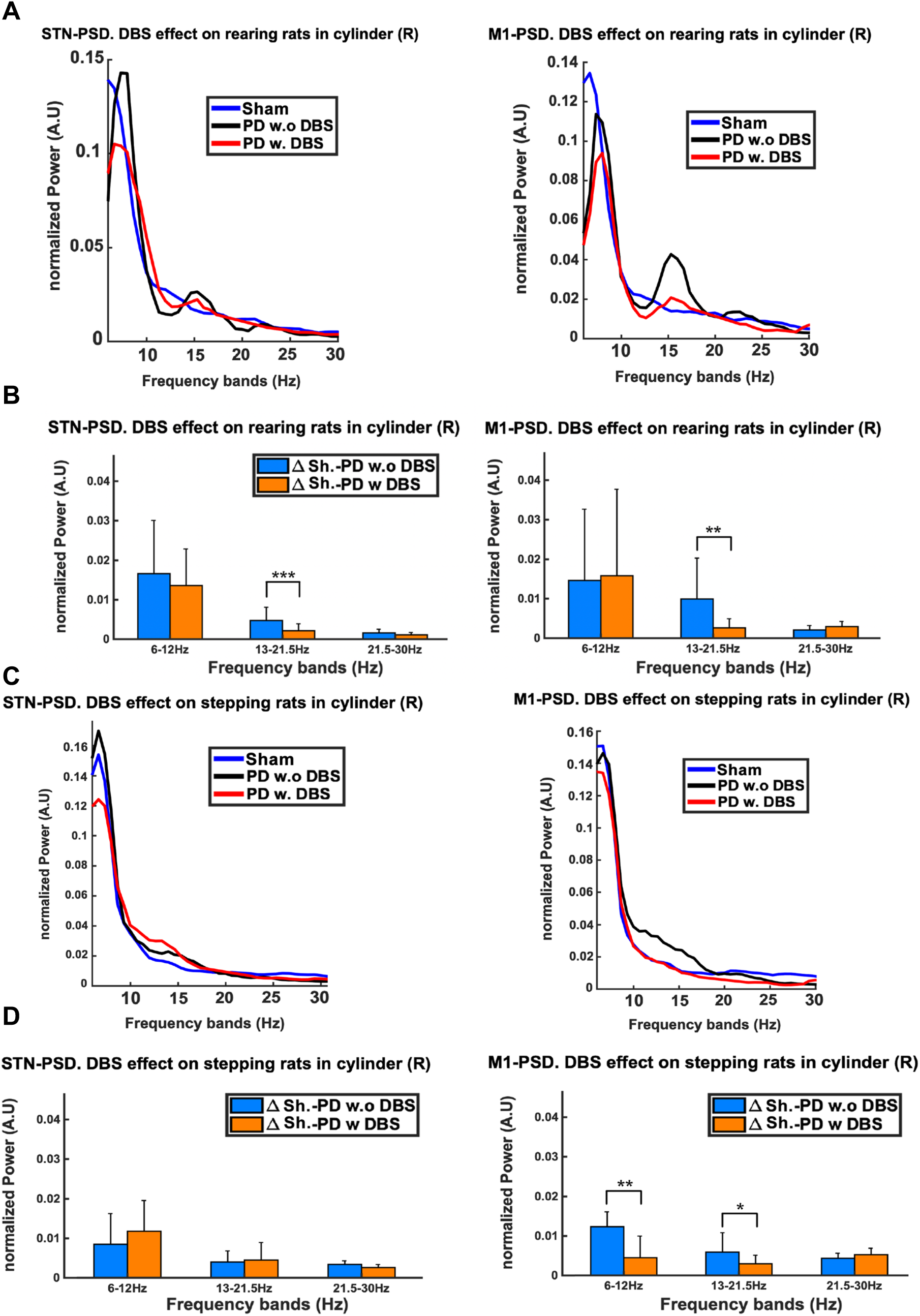
The excessive beta power of PD rats during stepping and rearing was compared between STN and M1 regions. A) Although excessive beta can be seen in both regions, the M1 exhibits a more pronounced excess during rearing as compared to the STN. C) The beta power excess during stepping can only been seen for M1 and not for the STN. We also observe that the administration of DBS lowers the beta excess power in both rearing and stepping episodes. B and D) Euclidean Distance (ED) was used to quantify the differences between the PSDs of PD DBS-ON vs. sham and PD DBS-OFF vs. sham rats. The low beta band shows significant decreases in the ED’s upon stimulation, which can be observed in both the STN and M1 during rearing, but only in M1 during stepping. Level of significance: (* p<0.05, ** p<0.01, *** p<0.001).

The PSD plots show how, in both rearing and stepping episodes, the excess in beta power of the DBS-OFF recordings from the lesioned regions is more prominent in M1 than in the STN. In addition, we observe how the application of DBS suppresses this excessive beta power in both regions during these episodes. The results of the ED analysis show that in rearing episodes (compared to no stimulation) DBS substantially lowers the ED’s in the low beta band for both the STN and M1. A similar analysis for stepping episodes shows a significant reduction in the ED’s for the low beta band only in M1 (not in the STN).

These results are very insightful as they tell us first, that the signals from M1 provide us with more clear and marked differences between electrophysiological signals coming from lesioned and healthy hemispheres, and are hence better suited for biomarker applications. These differences are also observed in more behaviors (i.e. both rearing and stepping) than for the STN recordings. Secondly, it quantitatively re-confirms the result from the previous section that the differences in the beta band power between lesioned and healthy hemisphere recordings become more pronounced as the animals’ activities become more complex and intensive. We also re-confirm the therapeutic effects of DBS, which returns the beta band power levels of PD rats close to healthy levels.

### D. The Low Beta Band is Related to Locomotion

To investigate the correlation between movement speed and spectral power in sham and PD rats, events involving locomotion were selected for the assessment of the OF test. For each locomotion speed in the range 1-16cm/s (1cm/s resolution), we analyzed LFP signals recorded from M1. In order to assess the effect of voluntary movement speed on the power in the different frequency bands, the average band power calculated over 5-second windows was determined at each speed and is illustrated in Figure 8. The violin plots on the right side of each graph represent the distribution of the power in each band. Additionally, the raster plots placed along the top of the figure indicate statistical significance for the difference in power between the sham and PD group at each speed.

**Figure 8.**
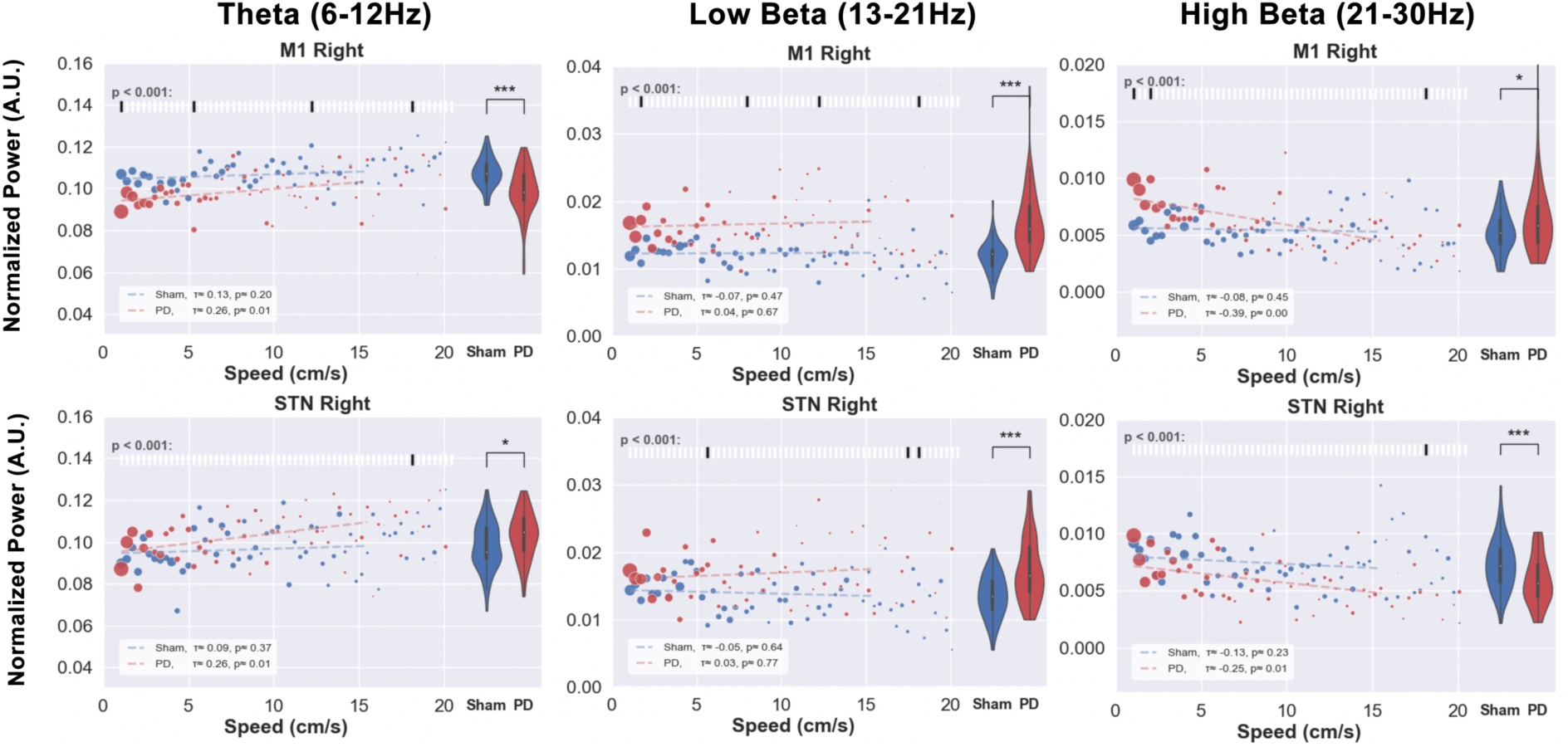
Speed-power relationship was analyzed in order to investigate the correlation between behavioral impairments and electrophysiological biomarkers. The locomotion speed of the animals was computed using the tracking data and the electrophysiological signals (M1 and STN) of the same time stamps were filtered in theta, low and high beta bands. The scatter plots above illustrate the median value of the band power at each specific speed (red for PD and blue for sham). The size of each point represents the relative amount of data per point. The violin plots on the right side of each figure represent the distribution of the powers over all the speeds. The significance test between PD and sham band powers at each speed was performed and is shown in the raster plot along the top of each subfigure. Low beta band power measured from the STN and M1 shows higher overall values in PD compared to sham rats, while the theta and high beta band powers measured from the STN and M1 exhibit contradictory patterns. No strong significance was shown in any of the bands among speed-power pairs. Level of significance: (* p<0.05, ** p<0.01, *** p<0.001).

Despite the weak differences between the individual speed-power pairs, the overall theta and low beta bands exhibited significant differences in power between the groups. Compared to sham group rats, PD group rats demonstrate lower theta and higher low beta powers. Additionally, Kendall’s rank correlation coefficient (τ) was computed to assess the linear correlation between speed and power band values, as shown in the white box in the bottom-left corner of each subfigure. For the sham group, no significant correlation between speed and power was found in any frequency band. The results for PD rats showed a small positive correlation in the theta band and a negative correlation in the high beta band.

The analysis performed on the STN recordings was in line with the results from M1 for the low beta band. However, for the theta and high beta bands, the STN and M1 showed differences in the relative power distributions of sham and PD animals: in the theta band, sham rats had higher powers overall in the M1 and lower in the STN. Similarly in the high beta band, powers were higher in M1 of PD rats, as compared to sham rats, and smaller in the STN.

The results of this experiment tell us that free movement speed does not influence the band powers significantly in neither sham nor parkinsonian rats. This indicates that the aforementioned beta band power biomarker is not suitable for revealing impediments in the movement speed of PD animals, impediments which are, however, clearly a symptom of the lesion as can be seen from the results in Figure 4. However, we reassuringly continue to see the excessive low beta power in PD rats, even during movement episodes. This power asymmetry between healthy and diseased animals, which has been observed in all experiments, could confirm the impact of frequency modulation in Parkinsonian motor impairments.

### E. DBS Restores Locomotion and Balance during Forced Movements

Both the cylinder and OF tests assess exploratory behavior by leaving space for the animal to start and stop movements freely. It was shown in previous experiments that PD animals tend to explore less than sham group animals. This motivated us to challenge their locomotion capability further by employing the rotarod test, in which the rats were forced to initiate movement and continue to walk to avoid falling. We measured the maximum speed and time that each rat was able to stay on the rotarod and compared results across groups (see Figure 9).

**Figure 9.**
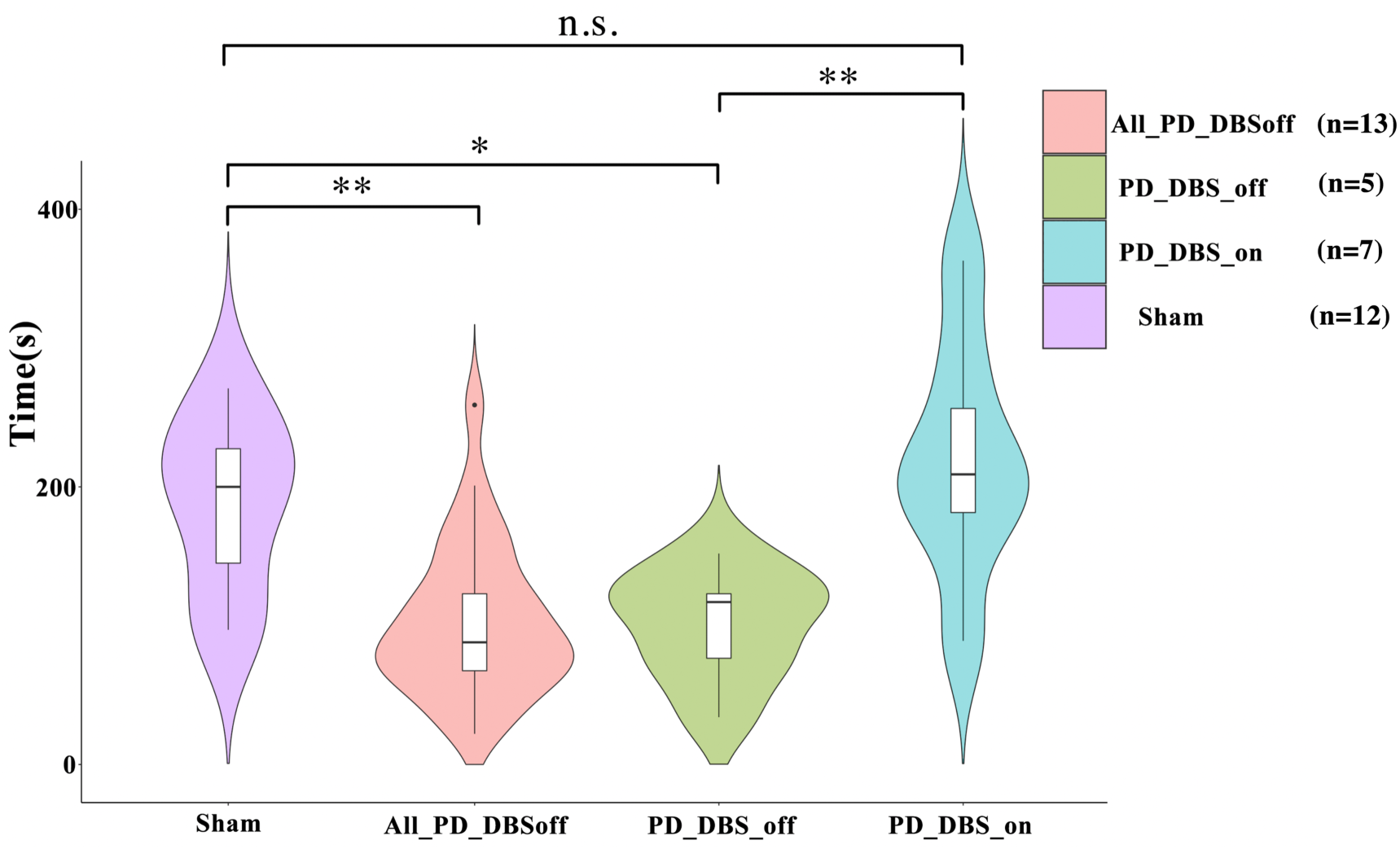
The rotarod test challenges the motor and balance capabilities of animals by forcing them to move at the speed of the elevated rotating rod they are placed on. All the animals underwent two test sessions while no DBS treatment was applied. A subgroup of PD animals performed a third test session while receiving DBS administration (n=7), whereas the rest of the PD group performed the third round while DBS was off (n=6). The maximum time and speed each animal could remain on the rotarod was recorded and compared across groups. The performance of the PD and sham group during the first two rounds were compared (sham vs. All_PD_DBSoff). The PD group remained a significantly shorter period of time on the rotarod compared to the sham group. As for the third round, the performance difference between the PD_DBS_on and sham animals was not significant. However, the performance of the PD_DBS_off subgroup was significantly worse compared to the sham and PD_DBS_on groups. Level of significance: (* p<0.05, ** p<0.01, *** p<0.001).

No significant difference was found between the first and the second round of trials in the sham group, meaning that we can compare the results of any of the two sham rounds to the other trials. Looking at the complete set of results, PD rats were found to stay a considerably shorter period of time on the rotarod compared to the sham group when no DBS was applied. However, remarkably, the administration of DBS greatly increases the time spent on the rotarod for PD rats (n=7) as compared to PD rats without DBS (n=6). Moreover, no significant differences were evident between the sham and PD-DBS group.

This experiment has shown us that the same conclusions we drew from the open field and cylinder tests remain valid in the realm of forced movements: lesioned animals have impaired movement capabilities compared to healthy animals and DBS is capable of restoring performance back to healthy levels. Here we have the added result that the lesion affects balance as well as movement abilities, and we continue to see that DBS is able to restore balance to its original state.

### F. Impairment of Movement Initiation in Hemi-Parkinsonian Rats is Accompanied by Alterations in Frequency Band Power

As for the OF test, a speed-power analysis was conducted for the rotarod test. Figure 10 shows how PD rats moved with higher speeds compared to the OF test. In all the selected bands, significant differences were exhibited between the speed-power pairs of PD and sham group animals (see raster plots), with powers being significantly lower for the PD rats in the theta band and higher in the other two bands compared to sham rats. Interestingly, the significance test between the powers at each speed showed significant differences in the low beta bands only during the early phase of the experiment, disappearing when the speed reaches 4 cm/s. The theta and high beta bands instead show strong differences at higher speeds. This dependence on speed of the significance in the difference between speed-power pairs between groups was not seen in the analysis of the OF test. Moreover, contrasting with the results from the OF test, the analysis of recordings from the STN showed patterns similar to those from M1.

**Figure 10.**
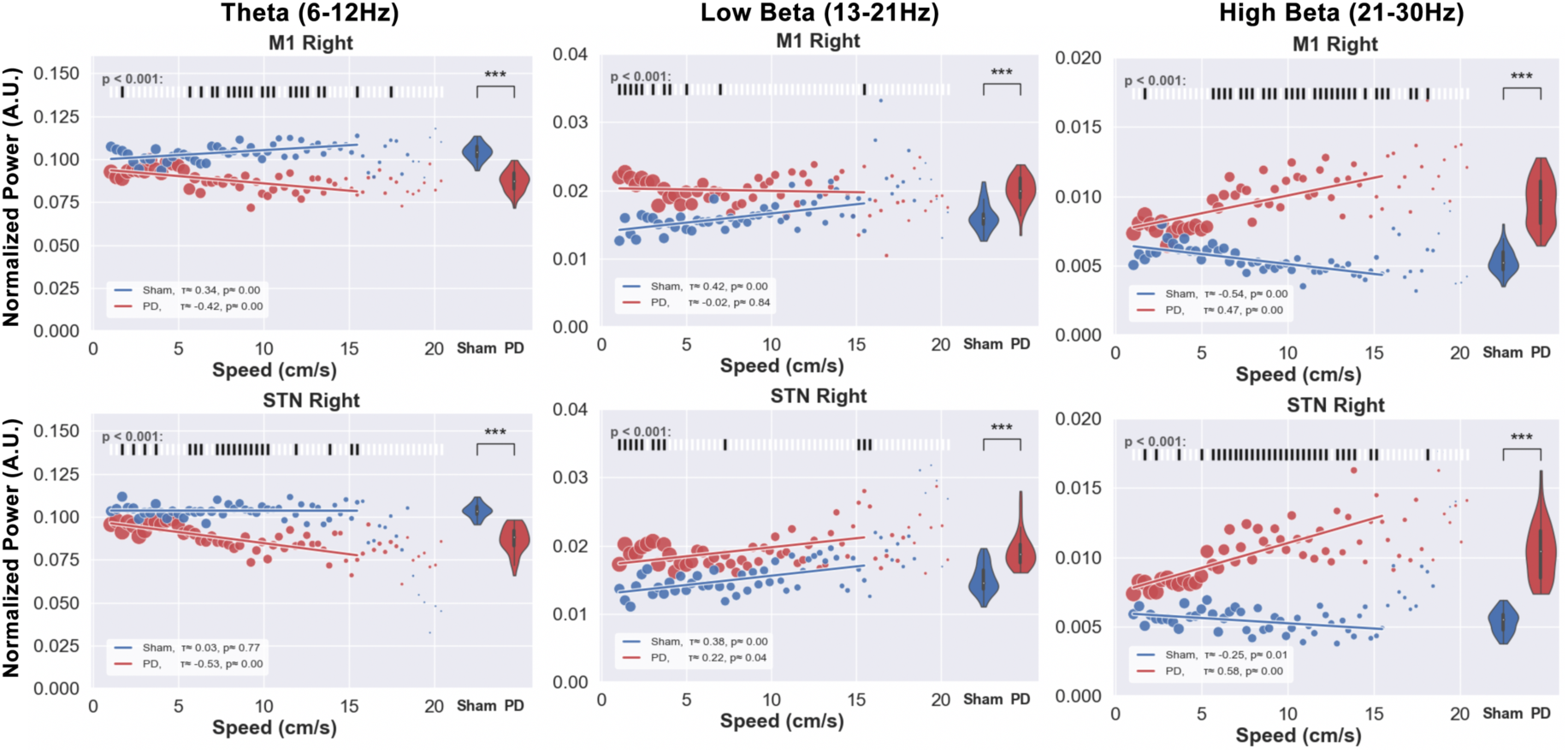
The relation between speed and band power in forced movements on the rotarod was investigated for PD and sham rats. The speeds of the rotarod as well as the electrophysiological signals (M1 and STN) of the same time stamps were plotted for theta, low and high beta bands. Each data point represents the median of the band power measured at each speed. The size of each point represents the relative amount of data per point. The distribution of the powers over all the speeds is represented by the violin plots on the right side of each subfigure, and the results from the significance test of the difference between the PD and sham band powers at each speed are illustrated in the raster plots above the figures. The theta powers measured from both the right STN and M1 regions over all the speeds (1-16 cm/s) are significantly lower in PD rats compared to the sham group, while both low and high beta powers showed higher values for PD compared to sham rats. Unlike the results from the OF test, the results from the STN are in line with the results from M1. Additionally, the low beta band shows stronger significant differences between PD and sham speed-power pairs only at lower speeds (<4 cm/s) while theta and high beta bands exhibit significant differences at higher speeds (> 4cm/s). Level of significance: (* p<0.05, ** p<0.01, *** p<0.001).

## Discussion

In the present study we have aimed to deepen our knowledge about the 6-OHDA PD rat model by conducting three behavioral-electrophysiological tests to evaluate the simulated motor symptoms, potential biomarkers and the efficacy of DBS at recovering motor impairments. The core pathology of PD is the progressive loss of dopaminergic neurons in the SNc (Agamanolis, 2016). This loss results in a shift of the brain’s electrophysiological profile as well as in distinctive motor symptoms. It is hence beneficial to investigate electrophysiological markers alongside behavioral features to characterize the effects of this disease, and this paper aims to provide such an understanding. Very limited research has been done in the discovery of electrophysiological biomarkers for this model of PD (Dorval et al., 2014; Anderson et al., 2015; Hoang et al., 2017) and these few studies focus mainly on the basal ganglia, without analyzing M1. Hence our results provide new perspectives on the potential of this model as a platform for the development and testing of closed-loop DBS.

Enhanced beta oscillatory activity throughout the cortico-thalamo-basal ganglia loop has been repeatedly reported in PD patients and pre-clinical animal models of the disease (Levy et al., 2002; Kühn et al., 2008; Dorval et al., 2010; Delaville et al., 2015; Brown, 2006). Studies show that it is mainly related to bradykinesia and akinesia symptoms and is thought to be an antikinetic feature of the disease, as dopamine replacement therapies (Heimer, Bar-Gad, Goldberg, & Bergman, 2002; Levy et al., 2002; Priori et al., 2004; Sharott et al., 2005; Williams, 2002) and DBS treatments (Wingeier et al., 2006) have been observed to lower its amplitude. Our results have shown that excessive beta power is exhibited in both the measured locations (STN and M1) of lesioned hemispheres, with its value being more prominent in M1.

The motor impairments observed in the model (less exploration, slower locomotion speeds, longer immobility) can be mostly linked to bradykinesia and akinesia, while no resting tremor was observed in any of the PD animals. This finding agrees with previous reports on the model (Asakawa et al., 2016). Moreover, we noticed that the beta power enhancement was not observed during inactive episodes, corroborating the similar results published by Degos et al., (2009). It has also been previously reported that the hemi-PD animal model expresses less exploratory behavior compared to healthy animals (Asakawa et al., 2016; Degos et al., 2009). Our experiments supported this finding as it was confirmed by the lesioned animals’ performances in the cylinder, OF and rotarod experiments. We further found that DBS treatment improves the performance of PD animals across all the tested experimental settings, suggesting an analogous DBS mechanism in rats to that of humans.

Among the behavioral features evaluated in the free movement experiments, rearing showed the most prominent elevation in the beta band power in lesioned hemispheres. The elevation covered the whole beta band and was detectable from the movement’s onset, whereas the prominence of the increased beta power at times just before and after the other free exploratory activities (e.g. stepping and walking) was not as strong. The need for averaging over large sets of data for the detection of excessive beta power in free stepping and walking could have lowered the temporal precision and hence the prominence of the measure. This inconvenience could greatly compromise this biomarker’s efficiency in closed-loop applications. Nevertheless, we promisingly observed that when the animal was forced to move on the rotarod, at all speeds the excessive beta power was detected much more easily than during free movements.

Our results on the change from significant excessive low-beta power at low speeds to excessive high-beta power as the speed surpasses approximately 4 cm/s highlights the importance of investigating the high and low beta bands individually. It has been shown before that the power in the high beta band is an important biomarker for cortico-subthalamic interactions affected by PD (Cao et al., 2019; Hirschmann et al., 2013; Lalo et al., 2008), while the low beta power correlates more with walking behaviors (Singh et al., 2013). Our results confirm this since excessive low beta was only detected during stepping behaviors in the cylinder and during walking behaviors in the OF.

Note that in all experiments the beta band was divided into an upper and lower portion because they reveal important features that are otherwise not visible. A good example is the strong positive correlation between the high beta power and speed in the rotarod test which would have been occluded by the weaker correlation of the low beta band, had the two been combined (Figure 10). Another important characteristic which would not have been observed is the divergence (with increasing speed) of the PD powers from the sham powers in the high beta band and the opposing convergence in the low beta band. This difference shows that, at low speeds, the low beta power might be a better indication of speed in PD rats than the high beta power, as it begins further away from the values for sham rats.

Valuable studies have attempted to describe behavioral tests in hemi-PD rodent models to quantitatively evaluate the severity of PD symptoms, to investigate the effects of novel therapeutic interventions and to gain insights into PD pathophysiology (Iancu, Mohapel, Brundin, & Paul, 2005; Sleeman, Boshoff, & Duty, 2012; Tan et al., 2015). However, like other neurotoxic models, the acute neurodegenerative property of the 6-OHDA model lacks the progressive, age-dependent effects of PD. Another problem is that characteristic biological traits such as the occurrence of Lewy bodies cannot be simulated by this model (Potashkin, Blume, & Runkle, 2011). We also observed that the 6-OHDA model does not simulate resting tremor, an important symptom of PD in humans. This explains why no salient features were detected in signals recorded during periods of inactivity. Moreover, the presence of some of the biomarkers found in this model remains to be confirmed in human PD patients, answering the question of the transferability of results from this rat model to humans. To-date, there are no behavioral models that can reproduce all of the symptoms that are commonly found in PD patients. Finally, the causal importance of the biomarkers needs to be established before they can be used to pattern stimulation in closed-loop DBS.

Across all the experiments and behaviors, the power in the high beta band was observed to be an important biomarker for PD as it showed differences between healthy and lesioned hemispheres and between PD and sham rats. This finding is not dissimilar from what we know about beta oscillations in humans, which have been previously proposed as a candidate variable for closed-loop control of DBS (Little & Brown, 2012). We have hence proved that this rat PD model not only simulates the motor symptoms precisely, but it also correctly models important electrophysiological characteristics of human PD.

Future work could be that of assessing the beta power of individual rats across several experiments such as free exploration, forced movement and drug treatment protocols to establish a pipeline in detecting the major and dominant frequencies within the beta band across different conditions. This would help develop an adaptive biomarker based on the animal’s state and hopefully prove beneficial for the treatment of PD patients.

